# Alterations in steady-state synchronisation between Acute and Chronic Tinnitus suggests reduction in both tinnitus and central gain

**DOI:** 10.1101/2025.08.28.672910

**Authors:** Abishek Umashankar, William Sedley, Kai Alter, Phillip E. Gander

## Abstract

The mechanism of tinnitus remains unclear as the generation and persistence of tinnitus is not widely understood. The acute tinnitus population serves as an ideal group for investigating the mechanisms behind the origin and persistence of tinnitus, from its initial onset to its subsequent chronification. One of the neural markers for measuring tinnitus such as neural synchrony or central gain is the Auditory Steady State Response (ASSR). The experiment was carried out on 39 participants with acute tinnitus (18 followed back six months post baseline), 30 with chronic tinnitus, and 27 controls. The results reveal that both cross sectionally and longitudinally, there were no significant differences in absolute amplitudes of ASSR across intensities and the slope of ASSR growth function between the groups. However, Controls tend to have an overall increase in ASSR amplitude when compared to Acute and Chronic Tinnitus which can be attributed to changes in neural synchronisation, attentional modulation, tinnitus-related distress, and diminished GABAergic inhibition coinciding with the presence of tinnitus. As there were no changes in the ASSR amplitude between Acute and Chronic Tinnitus or between Acute and Post Acute Tinnitus despite variations in distress and tinnitus loudness, we infer that auditory sensitivity/neural sensitvity is independent of the tinnitus.

## Introduction

Tinnitus has been found to originate most commonly because of hearing damage, which can be either an audiometrically detectable hearing loss or an undetectable kind of ‘hidden’ hearing loss such as cochlear synaptopathy (Kujawa & Liberman, 2015). In principle, for optimal functioning of the auditory system, each hierarchical level should have an input/output function that maintains a stable and optimum range of output values despite potentially large changes in the range of inputs. But, due to peripheral auditory insults, input to the central auditory system is reduced, thus altering the equilibrium of the input/output function. In order to maintain the input/output function, the brain typically changes the slope function by increasing the rate of excitatory to inhibitory firing to achieve homeostasis (Auerbach et al., 2014; Sedley et al., 2016). This alteration of slope in the input/output function is gain and helps achieve homeostatic plasticity (Sedley et al., 2016).

One of the objective markers of gain is the use of Auditory Steady State Response (ASSR) (Stapells et al., 1988). Amplitude Modulation (AM) results in neurons exhibiting a *mixed coding* response at the level of primary auditory cortex, which encompasses both phase locking to the modulation frequency, and an elevated firing rate in response to the unmodulated tone (Yin et al., 2011). The cortical source of ASSR at an AM of 40Hz is found to be at the level of primary auditory cortex with ASSR amplitudes increasing with stimulus intensity but reducing with increasing stimulus frequency (Ross et al., 2003; Wienbruch et al., 2006). Auditory steady-state responses (ASSRs) are also related to middle latency responses, which arise during the transition from early to late potentials, and even tapping into the inhibitory function (Yasoda-Mohan et al., 2024). The 40 Hz ASSR has been used in tinnitus studies as a potential measure of neuronal sensitivity in tinnitus. Most studies attribute increased ASSR in tinnitus patients when compared to non-tinnitus controls. A study conducted by Sadeghijam et al. (2023) examined the steady-state response across three groups: individuals with low distress tinnitus, individuals with high distress tinnitus, and non-tinnitus controls. Their findings indicated an elevation in ASSR for the low distress groups relative to the controls, whereas a decrease in ASSR was observed in the high distress group compared to the controls. Studies by Schlee (2006) and (Schlee et al., 2008) observed ASSR amplitude to be increased in tinnitus patients, and the response amplitude of the tinnitus frequency correlated with tinnitus loudness and distress, even after controlling for hearing loss. A recent study found out that there were enhanced 40 Hz and 80 Hz ASSR in tinnitus patients when compared to controls, attributing these differences to potential sensory impairments in higher-order auditory regions in individuals with idiopathic tinnitus. It is to be noted that these studies carry out measurements at a single intensity and infer changes in gain based on response amplitude (Ghasemahmad et al., 2024).

The slope typically dictates the degree of excess input/output function, which cannot be fully deduced from a single data point (intensity) measurement and requires at least 2 data points (intensities). Therefore, it is necessary to do an intensity dependence analysis (across different intensities) of the steady-state response, which will provide the slope function to facilitate the determination of central gain related to the ASSR. Another advantage of examining the intensity dependence of the ASSR is that it allows us to leverage each participant’s dynamic range, providing insight into how neural response amplitude changes in relation to perceived loudness growth.

The generation and persistence of tinnitus has been unclear and although there have been studies that talk about possible mechanisms of tinnitus generation and persistence, they are either from first principles, chronic tinnitus studies, or animal studies. There needs further research on acute tinnitus individuals as they are the ideal population to study how tinnitus generates and how does it transit from acute to chronic stages of tinnitus. According to the sensory precision model by Sedley et al. (2016), tinnitus arises from changes in the brain’s estimation of sensory precision following hearing loss. In the acute phase, reduced auditory input leads to a down-weighting of peripheral signals and an up-weighting of internal predictions, resulting in the emergence of tinnitus as a compensatory percept. Over time, if the altered sensory precision persists, the brain may recalibrate to treat the internally generated tinnitus signal as behaviourally relevant, leading to chronic tinnitus. This model suggests that persistent tinnitus reflects a maladaptive stabilization of heightened precision attributed to non-auditory predictive signals. Keeping this literature in mind, we hypothesize that central gain diminishes as a regression towards the mean from the Acute to the Chronic stage, leading us to predict a decline in ASSR slope and amplitude over time.

## Methods

### Participants

We studied 55 participants with Acute Tinnitus (which we defined as duration 3 days to 6 weeks, with one outlier having a duration of 12 weeks), 57 participants with Chronic Tinnitus (defined as duration more than 6 months), and 39 hearing-matched (to the acute tinnitus group) non-tinnitus controls. Participants were recruited through community advertising on Google Ads, and internally within Newcastle University’s research volunteer pool. The 55 Acute Tinnitus participants were invited for reassessment after a minimum of 6 months from tinnitus onset (which we took to indicate their chronic stage), and 26 of them volunteered for and completed this further testing. We refer to this as the ‘Post Acute’ group, to distinguish it from the group recruited during the chronic stage of tinnitus. The criteria for inclusion were as follows:

- Acute Tinnitus group: continuous non-pulsatile tinnitus present between 3 days and 6 weeks.
- Chronic Tinnitus group: continuous non-pulsatile tinnitus for more than 6 months
- Age 18 or over, with ability to provide informed consent.

Exclusion criteria:

- Meniere’s disease
- Epilepsy
- Middle ear pain or infections

Control participants were matched to the acute tinnitus group for age, sex, hearing, and stimulus frequency and presentation ear. Details are displayed in table 1.

**Table 1:**
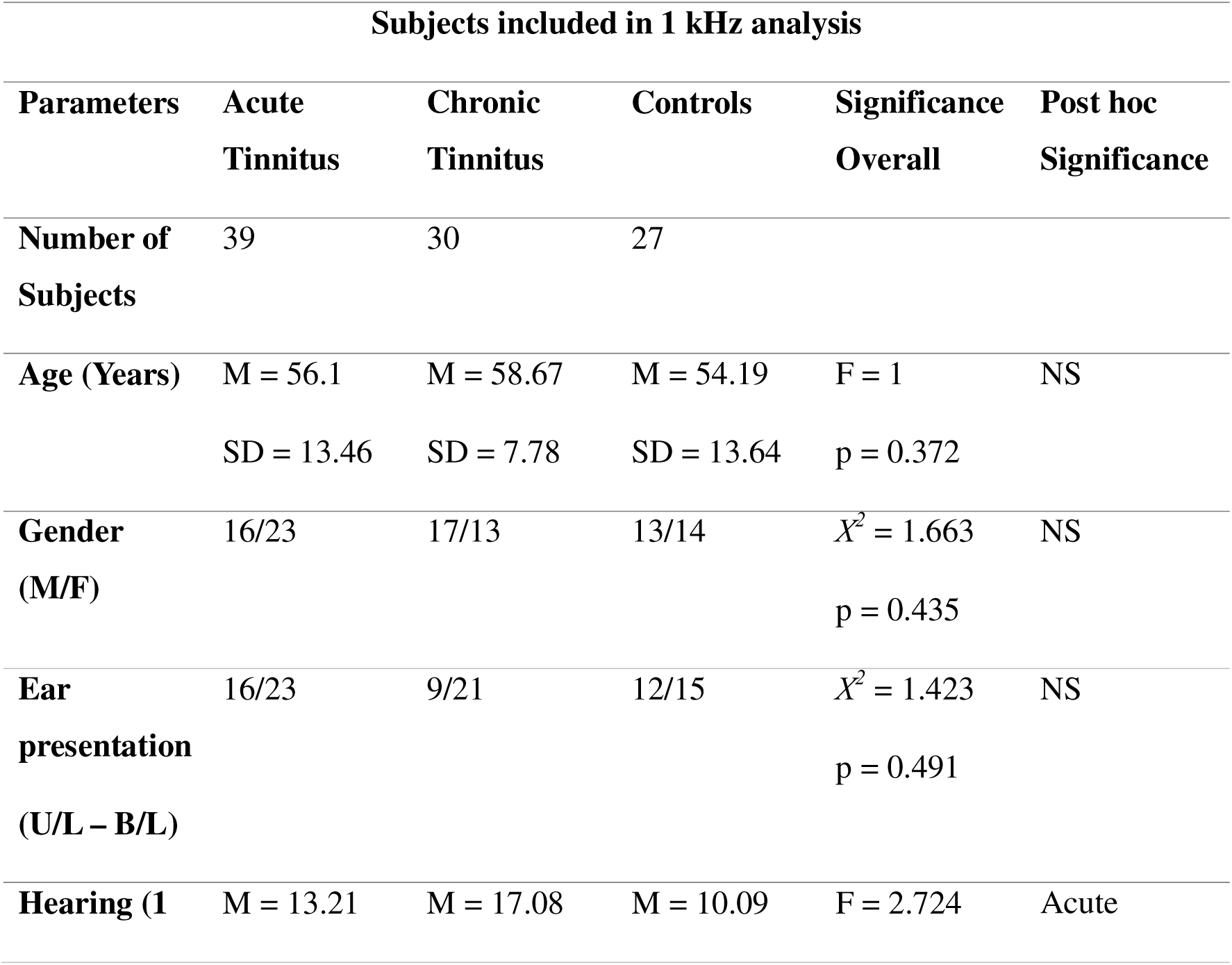
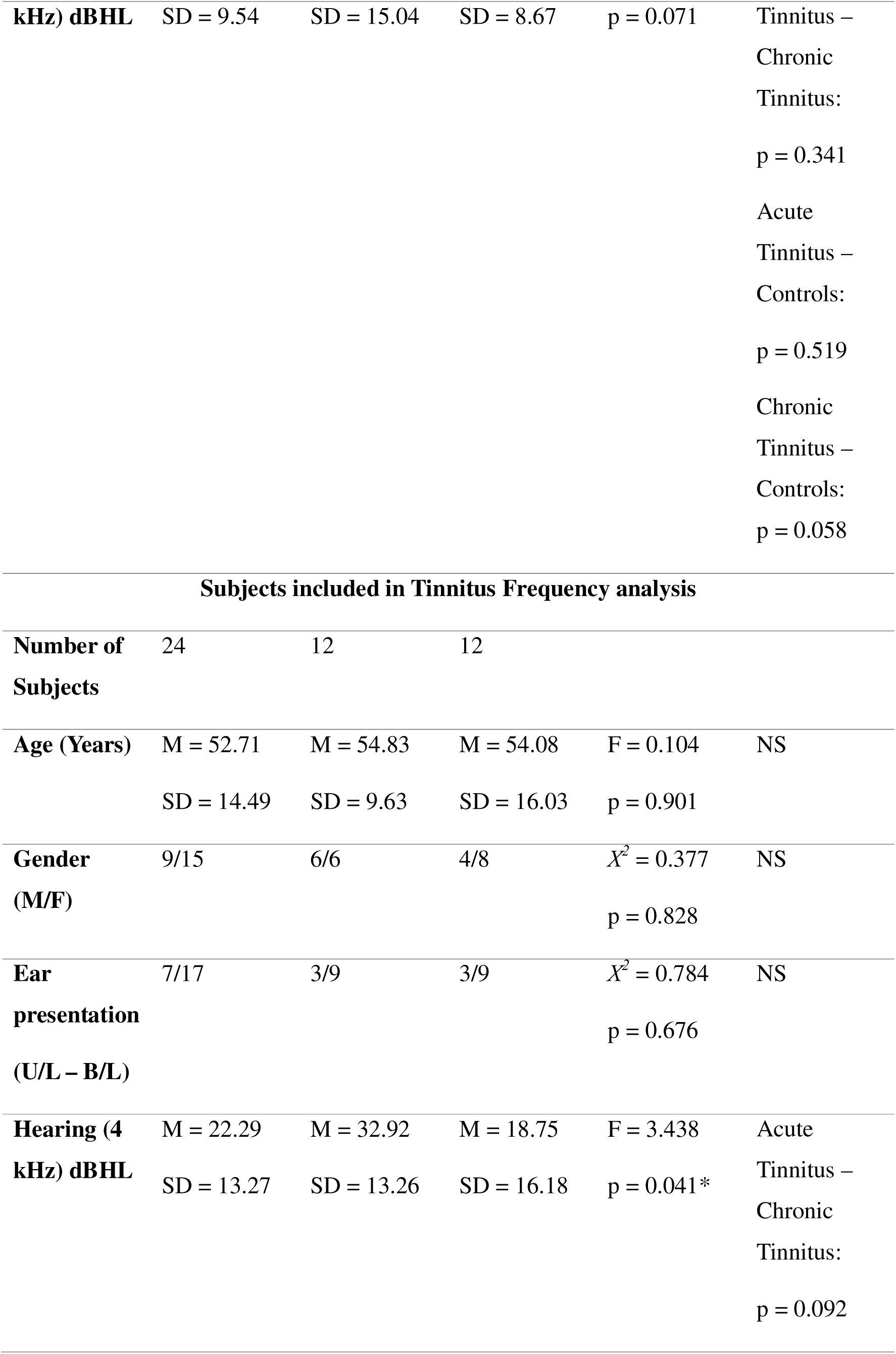
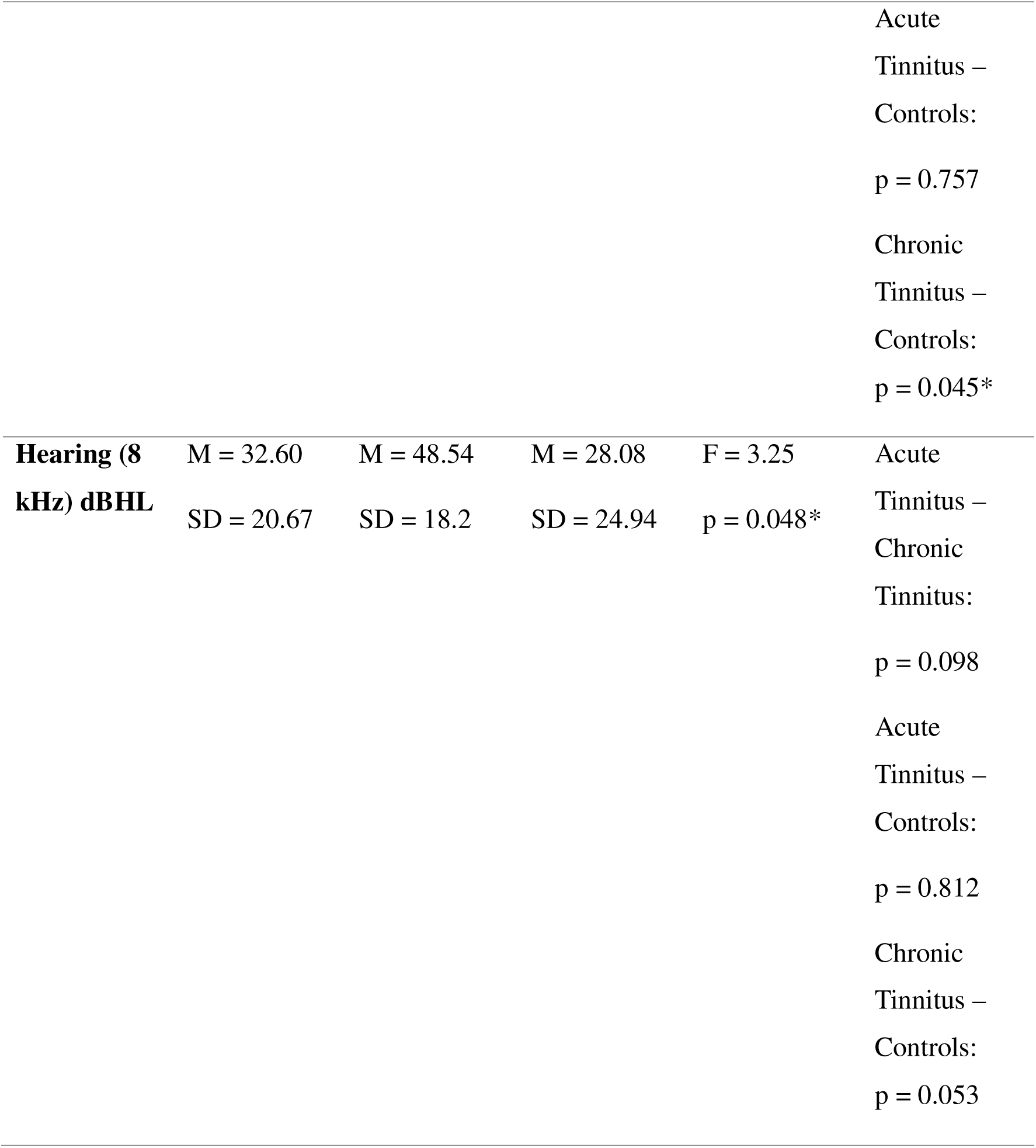
Participant demographics for each group. M denotes mean, SD denotes Standard Deviation, MD denotes Median, F statistic indicates a One Way ANOVA has been carried out, X^2^ indicates a chi square goodness of fit test, NS denotes No Significant difference, * indicates presence of statistical significance at 95% confidence interval. Hearing thresholds were calculated by averaging the left and right ear values in cases of bilateral tinnitus, whereas for unilateral tinnitus, the threshold was based on the ear with tinnitus.

| **Subjects included in 1 kHz analysis** |  |  |  |  |  |
| --- | --- | --- | --- | --- | --- |
| **Parameters** | **Acute**<br>**Tinnitus** | **Chronic**<br>**Tinnitus** | **Controls** | **Significance**<br>**Overall** | **Post hoc**<br>**Significance** |
| **Number of**<br>**Subjects** | 39 | 30 | 27 |  |  |
| **Age (Years)** | M = 56.1<br>SD = 13.46 | M = 58.67<br>SD = 7.78 | M = 54.19<br>SD = 13.64 | F = 1<br>p = 0.372 | NS |
| **Gender**<br>**(M/F)** | 16/23 | 17/13 | 13/14 | $X^2 = 1.663$<br>p = 0.435 | NS |
| **Ear**<br>**presentation**<br>**(U/L – B/L)** | 16/23 | 9/21 | 12/15 | $X^2 = 1.423$<br>p = 0.491 | NS |
| **Hearing (1** | M = 13.21 | M = 17.08 | M = 10.09 | F = 2.724 | Acute |
| **kHz) dBHL** | SD = 9.54 | SD = 15.04 | SD = 8.67 | p = 0.071 | Tinnitus –<br>Chronic<br>Tinnitus:<br>p = 0.341<br>Acute<br>Tinnitus –<br>Controls:<br>p = 0.519<br>Chronic<br>Tinnitus –<br>Controls:<br>p = 0.058 |
| **Subjects included in Tinnitus Frequency analysis** |  |  |  |  |  |
| **Number of**<br>**Subjects** | 24 | 12 | 12 |  |  |
| **Age (Years)** | M = 52.71<br>SD = 14.49 | M = 54.83<br>SD = 9.63 | M = 54.08<br>SD = 16.03 | F = 0.104<br>p = 0.901 | NS |
| **Gender**<br>**(M/F)** | 9/15 | 6/6 | 4/8 | $X^2 = 0.377$<br>p = 0.828 | NS |
| **Ear**<br>**presentation**<br>**(U/L – B/L)** | 7/17 | 3/9 | 3/9 | $X^2 = 0.784$<br>p = 0.676 | NS |
| **Hearing (4**<br>**kHz) dBHL** | M = 22.29<br>SD = 13.27 | M = 32.92<br>SD = 13.26 | M = 18.75<br>SD = 16.18 | F = 3.438<br>p = 0.041* | Acute<br>Tinnitus –<br>Chronic<br>Tinnitus:<br>p = 0.092 |

**Table 1: Participant demographics for each group.**
|  |  |  |  |  |  |
| --- | --- | --- | --- | --- | --- |
|  |  |  |  |  | Acute<br>Tinnitus –<br>Controls:<br>p = 0.757<br>Chronic<br>Tinnitus –<br>Controls:<br>p = 0.045* |
| **Hearing (8**<br>**kHz) dBHL** | M = 32.60<br>SD = 20.67 | M = 48.54<br>SD = 18.2 | M = 28.08<br>SD = 24.94 | F = 3.25<br>p = 0.048* | Acute<br>Tinnitus –<br>Chronic<br>Tinnitus:<br>p = 0.098<br>Acute<br>Tinnitus –<br>Controls:<br>p = 0.812<br>Chronic<br>Tinnitus –<br>Controls:<br>p = 0.053 |

### Audiological Assessment

The experimenter interviewed the subjects before the experiment, verifying a medical history consistent with subjective tinnitus and the absence of atypical symptoms or worries over an undetermined underlying cause. Before this, participants completed a concise, individualised pre-screening questionnaire that evaluated the inclusion and exclusion criteria. Participants submitted demographic data, including age and sex, alongside details regarding their tinnitus, including type (tonal, noise-like, or other), duration, affected ear(s), and a historical account of physical and mental health, particularly any otological disease. Four common, validated questionnaires—

- The Tinnitus Handicap Inventory (THI) (Newman et al., 1996).
- The Tinnitus Functional Index (TFI) (Meikle et al., 2012).
- The Hyperacusis Questionnaire (HQ) (Khalfa et al., 2002).
- The Inventory of Hyperacusis Symptoms (IHS) —were used to evaluate the impact and distress related to Tinnitus and any co-existing hyperacusis symptoms (Greenberg & Carlos, 2018).

Participants underwent a Pure Tone Audiometry (PTA) test to establish hearing thresholds at octave frequencies ranging between 250 Hz up to 8 kHz. Thresholds were estimated based on the initial presented level at either 40 dBHL or above depending on the participant’s residual hearing. If perceived audible, the stimulus was continuously reduced by 15 dBHL until the participant did not perceive it as audible. From the first reversal, a 5 dB up, 10 dB down staircase procedure was then used until the final threshold was established by two positive responses out of three trials.

### Tinnitometry

The frequency and loudness of tinnitus were assessed using a tinnitometry approach for patients with acute or Chronic Tinnitus. In instances of unilateral tinnitus, matching stimuli were delivered contralaterally, bilateral tinnitus, they were administered bilaterally. Participants initially matched the loudness of their tinnitus, followed by matching its frequency, beginning at a reference level of 6 kHz. In bilateral tinnitus cases, supplementary modifications were implemented to equilibrate the two ears when tinnitus presented asymmetrically. A procedure was implemented in which the loudness and then frequency were sequentially and iteratively modified until no further adjustments were deemed required by the participant to provide an optimal match. The technique was conducted over three trials, and the average of these trials was determined as the final intensity and frequency of the tinnitus. The stimulus employed for this method was either a pure tone or narrow band noise, contingent upon the subjective characteristics of the tinnitus.

### ASSR Stimuli

The ASSR stimuli comprised 15 stimuli per frequency (5 stimuli for low, medium and high intensities respectively) within a total experiment duration of 15 minutes. These stimuli were Amplitude Modulated sounds with a modulation frequency of 40Hz and a modulation depth of 100% at 1 kHz and the participant’s matched tinnitus frequency, each lasting 30 seconds, with an interstimulus interval of 2 seconds. In the Control group, tinnitus frequencies were not self-reported but were assigned based on age-, sex-, and hearing-matched counterparts from the Acute Tinnitus group. This approach allowed for consistent frequency-based comparisons across groups For every frequency, three different intensities were presented. Stimuli were presented in one block, with intensities and frequencies fully randomised within that block. To account for hearing loss, loudness recruitment and hyperacusis, whilst keeping the stimuli clearly audible yet comfortable, stimulus intensities were presented in accordance with each participant’s dynamic range, i.e. the range between their hearing threshold and uncomfortable loudness level, which were established for the specific experimental stimuli at both 1 kHz and the tinnitus frequency. Hearing thresholds were established using an ascending-descending run of dB steps until at least 50% positive responses were obtained (i.e. 2 correct responses out of trials). The uncomfortable loudness level was established by presenting an ascending run of 3 dB step size from the threshold until the participant just perceived the tone to be uncomfortably loud or showed signs of discomfort (‘Recommended Procedure Determination of uncomfortable loudness levels’, 2022). Dynamic range was defined as uncomfortable loudness level (ULL) minus hearing threshold. Stimulus intensities were determined as follows:

a. Low Intensity: hearing threshold plus 60% of Dynamic Range
b. High Intensity: ULL – 10 dB
c. Medium Intensity: Mean of Low and High Intensity

For those with unilateral tinnitus, stimuli were only presented in the tinnitus ear; for those with bilateral tinnitus, they were presented in both ears. Participants with Acute Tinnitus were matched individually with controls of the same age, sex, and hearing. Control participants were presented with stimuli that matched the frequency and presentation ear(s) of their matched tinnitus participant but were individualised in intensity to the control participant based on the same dynamic range procedure. Figure 1 depicts the ear and presentation levels and table 1 shows the ear and stimulus presentations for each group.

**Figure 1:**
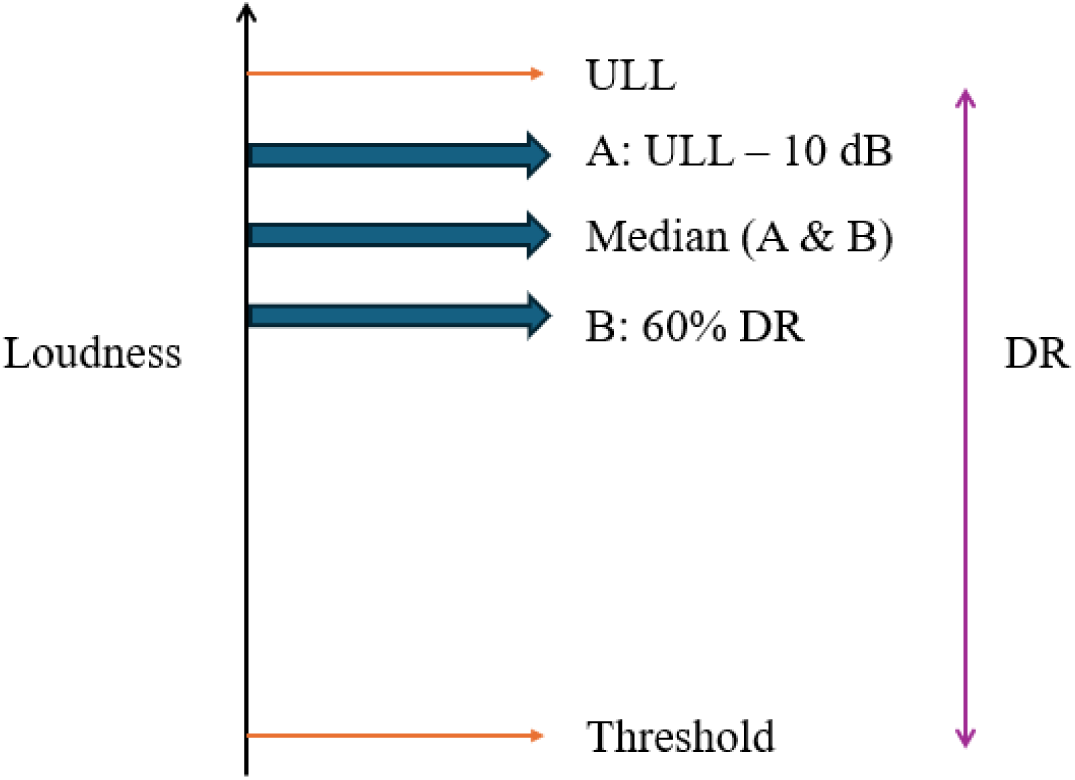
Stimulus presentation levels across 3 intensities based on tailored dynamic range for each subject *ULL: Uncomfortable Loudness level, DR: Dynamic Range*

### ASSR Recording

EEG was recorded using a 64-channel Active Two system (Biosemi) in a soundproof room. Electrode offset was kept at the manufacturer’s recommended limits of ±10mv with a sampling rate of 256Hz.

The experiment was a passive task, during which the participants watched a silent subtitled movie of their choosing. All stimuli were generated and presented using Matlab version R2019a, using the Psychtoolbox toolbox (Brainard, 1997; Pelli, 1997; Kleiner, Brainard, Pelli 2007).

### ASSR Pre Processing

Data were processed in MATLAB, version R2019a, using the EEGlab toolbox (Delorme A & Makeig S, 2004) and customised code. Data were re-referenced to the P9/P10 (Linked mastoids) montage. The data were further segmented into 30 seconds epoch around each trigger which was followed by a baseline correction. A frequency analysis was conducted utilising a Fast Fourier Transform (FFT) to transfer the time-domain data into the frequency domain, facilitating the identification and extraction of spectrum peaks, notably at 40Hz.

### Data Analysis

After pre-processing, we used customised code to perform automatic peak detection at 40Hz and signal to noise ratio calculation to determine the ASSR amplitude and signal to noise ratio. We did this using data from FCz to measure the peak amplitude for ASSR at 40Hz for both 1 kHz and tinnitus frequency. Similar analysis was done for noise floor estimation, where the FFT amplitudes were averaged across the neighbouring frequencies between 38 and 39, and between 41 and 42, Hz of modulation rate. The presence of a meaningful ASSR was ascertained via visual evaluation of the response at 40Hz and objectively through F-statistics (Korczak et al., 2012), which analysed the variance of peak amplitude and noise floor values; a meaningful response was judged to be present only if the ASSR response was significantly larger than the noise floor values based on a value of p < 0.05. Each subject had amplitude and noise floor calculated for 3 intensities across 2 frequencies of 1 kHz and tinnitus frequency. From these, the ASSR slope function at each frequency was calculated as the quotient of the Amplitude Dynamic Range (ADR: relative difference of the high and low ASSR amplitude) and Stimulus Dynamic Range (SDR: relative difference of the high and low stimulus intensity).

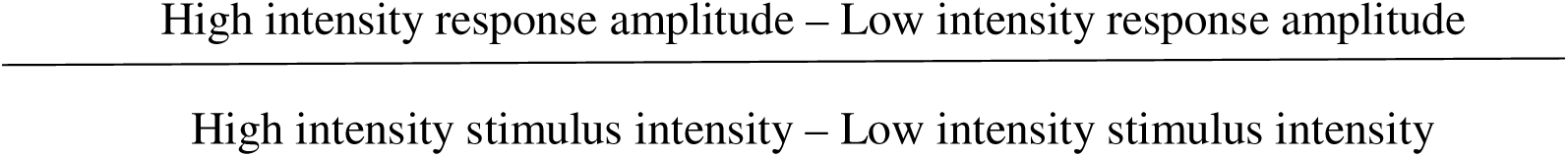

Units of the ASSR slope are thus nV/dB. Note that the mid-intensity stimuli were not used for analysis but were judged important to include in the paradigm for their influence on the overall statistics of presented stimuli, and adaptation occurring in the auditory system. Figure 2 depicts a sample ASSR response along with its noise floor.

**Figure 2:**
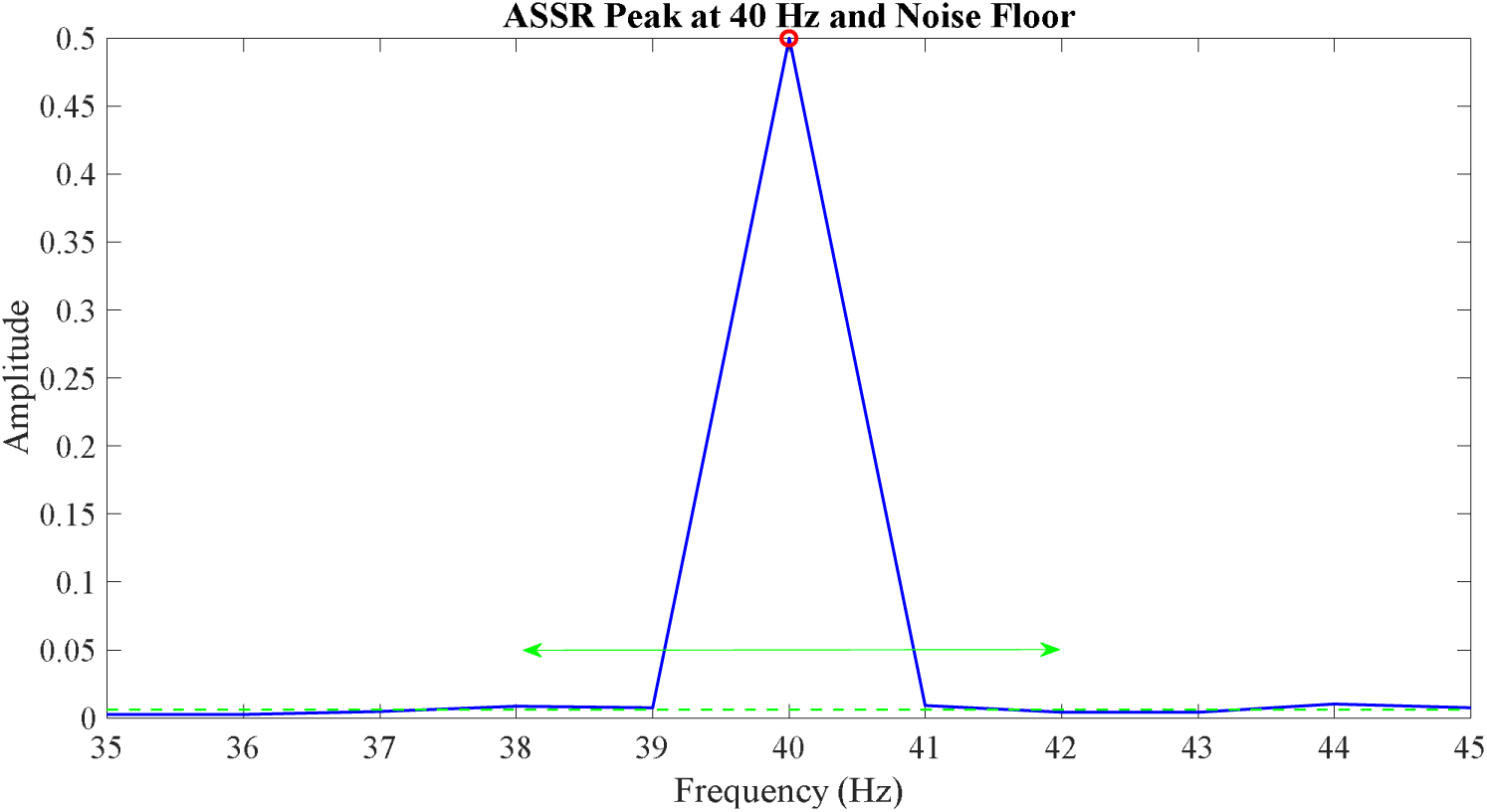
Example of an ASSR response at 40Hz along with its noise floor. The blue peaked line denotes the peak ASSR at 40Hz, while the green dotted lines indicate the noise floor across various modulation frequencies. The red circle denotes the peak amplitude of ASSR at 40 Hz, while the green double arrow represents the noise floor between 38 Hz and 42 Hz that is considered for statistical analysis. The F statistic will lie between the amplitude values indicated by the red circle and those of the noise floor (green dotted lines).

Statistical analysis was performed using the Statistical Package for Social Science (SPSS). Based on the Shapiro-Wilk test of normality, the data were normally distributed across the three groups for ASSR amplitude, and hence parametric statistics were used. However, the data were not normally distributed for ASSR slope at tinnitus frequency, and hence a non-parametric statistic was used for slope. The statistical analysis was done in two stages.

To examine the effect of stimulus intensity and frequency on ASSR amplitude across groups, we conducted a three-way factorial Analysis of Variance (ANOVA) with group (Acute, Chronic, and Controls) as the between-subject factor, and intensity (low, medium, high) and frequency (1 kHz and tinnitus frequency) as within-subject factors. The dependent variable was the stimulus presentation level. There there was no interaction effect between Group, frequency, and intensity (F(4,574) = 0.170, p = 0.954) indicating that there were no group differences in stimulus presentation and that stimulus presentation levels did not impact ASSR amplitude. To examine the ASSR amplitude across groups, we conducted a three-way factorial ANOVA with Group (Acute, Chronic, and Controls) as the between-subject factor, and Intensity (low, medium, high) and Frequency (1 kHz and tinnitus frequency) as within-subject factors. The dependent variable was ASSR amplitude. Main effects and interactions were tested, and when significant, post-hoc analyses were carried out using Tukey’s Honestly Significant Difference (HSD) test and Bonferroni-adjusted comparisons to account for multiple testing.

For the longitudinal comparison, there was no interaction effect for group, frequency, and intensity on the stimulus presentation levels (F(2,10) = 0.059, p = 0.943) and hence stimulus presentation levels did not imact ASSR amplitude. A three-way repeated measures ANOVA was conducted to examine the effects of group, frequency, and intensity on the ASSR amplitude. Assumptions of normality and sphericity were assessed prior to analysis.

Mauchly’s Test of Sphericity if significant for the three-way interaction, the degrees of freedom will be adjusted using the Greenhouse-Geisser correction. Where significant main effects or interactions were found, post hoc pairwise comparisons were conducted using the Bonferroni correction to control for multiple comparisons.

As a second analysis, the ASSR slope values for each frequency were compared between the groups. Since there existed sample size inequality between the stimulus frequency groups (table 1), statistical analysis was conducted independently for each frequency instead of a factorial analysis with group and frequency. Significant differences across the groups for 1 kHz were determined using a one-way ANOVA (Analysis of Variance), and for the tinnitus frequency groups, the Non-parametric Kruskal Wallis test was used. To compare the groups between the Acute phase and the follow-up Post Acute phase for both frequencies, a paired t test was employed. Figure 2 is an illustration that presents a sample waveform from a single subject.

**Figure.2:**
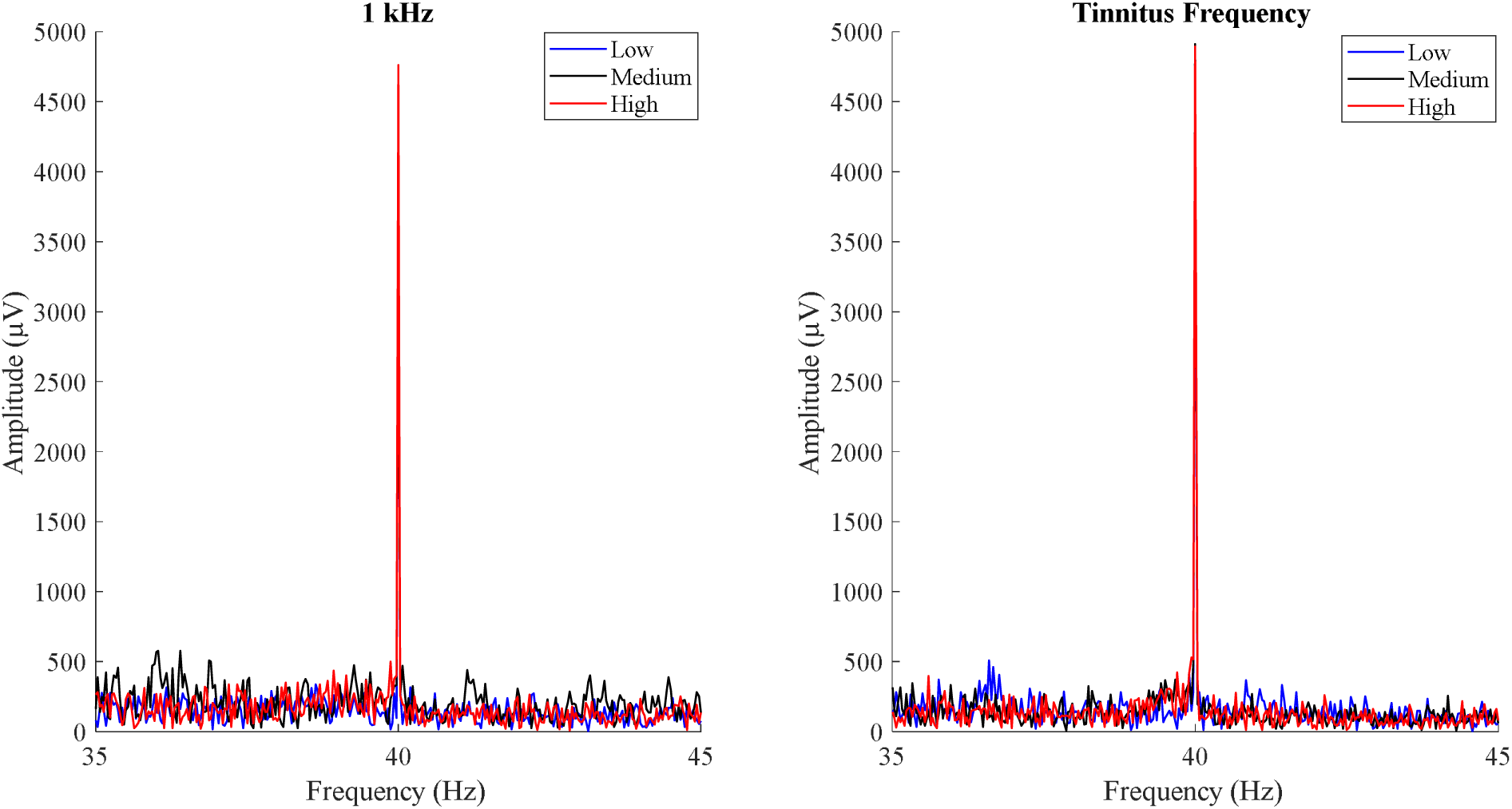
Example of an ASSR waveform for a subject. The waveforms are recorded across different intensities and frequencies at a 40ϑHz modulation rate, with the noise floor shown at neighbouring frequencies

## Results

The final demographic of participants following data quality-based rejection for ASSR slope was as follows: Acute Tinnitus (1 kHz – 39, Tinnitus Frequency – 24), Chronic Tinnitus (1 kHz – 30, Tinnitus Frequency – 12), and Controls (1 kHz – 27, Tinnitus Frequency – 12).

Participants were rejected based on their overall ERP quality (both visual inspection and F statistic) due to lower signal to noise ratio for the ASSR. Eighteen Acute Tinnitus returned six months post their baseline for their re-evaluation marking their onset of their chronic stage of tinnitus (Post Acute Tinnitus). Such exclusions were performed with the researcher blind to the group to which each participant belonged.

This section is divided into six sections: 1) demographics and hearing between the groups; 2) differences in tinnitus handicap between the groups; 3) differences in hyperacusis scores between the groups; 4) differences in ASSR amplitudes between the groups; 5) differences in ASSR slope between the groups; 6) summary of results. Different statistics were run for each frequency due to the variations in sample sizes between the tinnitus frequency and 1 kHz, and these will be discussed individually.

### Demographics and Hearing

Participant groups were attempted to be matched for age, sex, hearing, and mode of stimulus presentation. There were no significant differences across the groups for age, sex, and stimulus ear presentation for both the samples at 1 kHz and tinnitus frequency. Using the 1 kHz, 4 kHz, and 8 kHz hearing thresholds, an attempt was made to pair the groups. However, non-significant trend was present at 1 kHz (F(2,93) = 2.724, p = 0.0071) and a significant difference at 4 kHz (F(2,50) = 3.428 p = 0.041), and 8 kHz (F(2,46) = 3.25, p = 0.048), with all cases being that Chronic Tinnitus having higher thresholds than Acute Tinnitus and Controls (Table 1). After excluding certain subjects from each analysis, for data quality purposes, the composition of the groups including 1 kHz and tinnitus frequency analyses were slightly different, and we report each separately below. For further details please refer to table 1.

### Symptom Scores for Tinnitus Distress

With respect to symptom scores for the individuals included in 1 kHz cross-sectional comparisons between the Acute and Chronic Tinnitus for THI and TFI yielded significant differences between them, with Chronic Tinnitus having higher levels of distress than Acute Tinnitus (refer table 2). Comparison between the Acute and Post Acute Tinnitus group revealed a statistically significant difference with reduction over time in TFI scores with Acute Tinnitus having higher TFI scores when compared to Post Acute Tinnitus. No differences were noted between Acute and Post Acute Tinnitus for THI.

**Table 2:** Participant’s tinnitus characteristics for each group. M denotes mean, SD denotes Standard Deviation, MD denotes Median, F statistic indicates a One Way ANOVA has been carried out, t statistic indicates an independent t test done, z statistic indicates a non-parametric Man-Whitney u test has been carried out, N/A-Not Applicable, NS denotes No Significance. * indicates statistically significant difference at 95% confidence interval, THI - Tinnitus Handicap Inventory, TFI – Tinnitus Functional Index, HQ – Hyperacusis Questionnaire, IHS – Inventory of Hyperacusis Symptoms

| **1 kHz** |  |  |  |  |  |
| --- | --- | --- | --- | --- | --- |
| **Parameters** | **Acute**<br>**Tinnitus** | **Chronic**<br>**Tinnitus** | **Controls** | **Significance**<br>**Overall** | **Post hoc**<br>**Significance** |
| **Number of**<br>**Subjects** | 39 | 30 | 27 |  |  |
| **Tinnitus**<br>**Pitch (Hz)** | M =<br>5709.02<br>SD =<br>2064.88 | M =<br>6775.1<br>SD =<br>1410.25 | M =<br>5928.56<br>SD =<br>2251.04 | F = 2.703<br>p = 0.072 | NS |
| **Tinnitus**<br>**Loudness**<br>**(dBSL)** | M = 16.31<br>MD =<br>14.5<br>SD =<br>15.73 | M =<br>6.46<br>MD =<br>6.17<br>SD =<br>10.05 | N/A | z = 2.622<br>p = 0.009* |  |
| **Number of**<br>**Subjects** | 38 | 24 | 24 |  |  |
| **THI** | M = 30.58<br>SD =<br>21.04 | M = 44.33<br>SD =<br>20.68 | N/A | t = - 2.524<br>p = 0.014* |  |
| **TFI** | M = 95.6<br>SD = | M =<br>131.08 | N/A | t = -2.787<br>p = 0.004* |  |
|  | 45.79 | SD = |  |  |  |
|  |  | 53.35 |  |  |  |
| **HQ** | M = 12.79 | M = 17.54 | M = 8.13 | F = 7.597 | Acute Tinnitus– |
|  | SD = 7.61 | SD = | SD = 5.62 | p < 0.001* | Control: |
|  |  | 11.31 |  |  | p = 0.081 |
|  |  |  |  |  | Acute Tinnitus– |
|  |  |  |  |  | Control: |
|  |  |  |  |  | p = 0.088 |
|  |  |  |  |  | Chronic Tinnitus |
|  |  |  |  |  | – Control: |
|  |  |  |  |  | p < 0.001* |
| **IHS** | M = 41.18 | M = 52.16 | M = 31.79 | F = 13.516 | Acute Tinnitus– |
|  | SD = | SD = | SD = 7.58 | p < 0.001* | Chronic |
|  | 12.98 | 18.39 |  |  | Tinnitus: |
|  |  |  |  |  | p = 0.007* |
|  |  |  |  |  | Acute Tinnitus– |
|  |  |  |  |  | Control: |
|  |  |  |  |  | p = 0.026* |
|  |  |  |  |  | Chronic Tinnitus |
|  |  |  |  |  | – Control: |
|  |  |  |  |  | p < 0.001* |
| **Tinnitus Frequency** |  |  |  |  |  |
| **Number of** | 24 | 12 | 12 |  |  |
| **Subjects** |  |  |  |  |  |
| **Tinnitus** | M = | M = | M = | F = 3.142 | NS |
| **Pitch (Hz)** | 5051.43 | 6686.51 | 4577.1 | p = 0.052 |  |
|  | SD =<br>2266.91 | SD =<br>2144.69 | SD =<br>2159.78 |  |  |
| **Tinnitus** | M = 17.25 | M = 11.35 | N/A | z = 0.689 |  |
| **Loudness**<br>**(dBSL)** | MD =<br>11.25 | MD =<br>7.83 |  | p = 0.491 |  |
|  | SD =<br>18.04 | SD =<br>13.06 |  |  |  |
| **Number of**<br>**Subjects** | 24 | 11 | 10 |  |  |
| **THI** | M = 36.25 | M = 48.91 | N/A | t = -1.535 |  |
|  | SD = 24 | SD =<br>21.81 |  | p = 0.134 |  |
| **TFI** | M =<br>102.46 | M =<br>148.36 | N/A | t = -2.486 |  |
|  | SD =<br>46.23 | SD =<br>59.74 |  | p = 0.018* |  |
| **HQ** | M = 13.25 | M = 20.27 | M = 5.3 | F = 8.369 | Acute Tinnitus– |
|  | SD = 7.42 | SD =<br>12.41 | SD = 3.95 | p < 0.001* | Chronic<br>Tinnitus:<br>p = 0.066 |
|  |  |  |  |  | Acute Tinnitus–<br>Controls:<br>p = 0.04* |
|  |  |  |  |  | Chronic Tinnitus<br>– Control:<br>p < 0.001* |
| **IHS** | M = 43.12 | M = 58.27 | M = 28.8 | F = 10.28 | Acute Tinnitus– |
|  | SD = | SD = | SD = 3.01 | p < 0.001* | Chronic |
|  | 15.75 | 18.81 |  |  | Tinnitus: |
|  |  |  |  |  | p = 0.02* |
|  |  |  |  |  | Acute Tinnitus– |
|  |  |  |  |  | Controls: |
|  |  |  |  |  | p = 0.037* |
|  |  |  |  |  | Chronic Tinnitus |
|  |  |  |  |  | – Control: |
|  |  |  |  |  | p < 0.001* |

Comparable results were achieved when the identical comparisons were reiterated solely for the subjects whose data were utilised for the examination of reactions to the tinnitus frequency stimuli. There were no differences between the Acute and Chronic Tinnitus on THI, but differences were noted for TFI with Chronic Tinnitus having greater distress on TFI than Acute Tinnitus (refer table 3). Pairwise comparison yielded non-significant trend to significant differences between Acute and Post Acute Tinnitus group with Acute Tinnitus having higher THI and TFI scores when compared to Post Acute Tinnitus.

**Table 3:** Longitudinal changes between Acute and Post Acute Tinnitus. The t test denotes a paired wise comparison between the Acute and Post Acute Tinnitus group and z test indicates the non-parametric Wilcoxon signed rank test. * indicates presence of statistically significant difference at 95% confidence interval, THI - Tinnitus Handicap Inventory, TFI – Tinnitus Functional Index, HQ – Hyperacusis Questionnaire, IHS – Inventory of Hyperacusis Symptoms. One person did not complete the questionnaire and hence the sample size have been reduced in both 1 kHz and tinnitus frequency analysis.

| **1 kHz** |  |  |  |
| --- | --- | --- | --- |
| **Parameters** | **Acute**<br>**Tinnitus** | **Post Acute**<br>**Tinnitus** | **Significance**<br>**Overall** |
| **Number of**<br>**Subjects** | 18 | 18 |  |
| **Tinnitus**<br>**Frequency**<br>**(Hz)** | M = 5870.99<br>SD =<br>1791.45 | M = 6603.1<br>SD =<br>1626.18 | t = 1.256<br>p = 0.226 |
| **Tinnitus Loudness (dBSL)** | M = 13.99<br>MD = 11.25<br>SD = 10.98 | M = 9.08<br>MD = 5.33<br>SD = 10.11 | t = -2.266<br>p = 0.023* |
| **Number of Subjects** | 16 | 16 |  |
| **THI** | M = 31.86<br>SD = 20.3 | M = 25.5<br>SD = 27.15 | t = -1.626<br>p = 0.125 |
| **TFI** | M = 101.19,<br>SD = 43.94 | M = 75.81,<br>SD = 53.67 | t = -3.418<br>p = 0.004* |
| **HQ** | M = 12.75<br>SD = 8.41 | M = 13.13<br>SD = 8.94 | t = 0.816<br>p = 0.237 |
| **IHS** | M = 43.69<br>SD = 15.64 | M = 43.13<br>SD = 18.51 | t = -0.183<br>p = 0.857 |
| **Tinnitus Frequency** |  |  |  |
| **Number of Subjects** | 9 | 9 |  |
| **Tinnitus** | M = 5995 | M = 6431.42 | z = 0 |
| **Frequency** | MD = | MD = | p = 1 |
| **(Hz)** | 6081.49 | 6507.72 |  |
|  | SD = | SD = |  |
|  | 2409.61 | 1482.69 |  |
| **Tinnitus** | M = 18 | M = 14.037 | z = -0.71 |
| **Loudness** | MD = 9 | MD = 11 | p = 0.359 |
| **(dBSL)** | SD = 10.16 | SD = 9.98 |  |
| **Number of** | 8 | 8 |  |
| **Subjects** |  |  |  |
| **THI** | M = 33 | M = 25 | z = -1.759 |
|  | MD = 30 | MD = 17 | p = 0.079 |
|  | SD = 24.02 | SD = 29.02 |  |
| **TFI** | M = 102.12 | M = 71 | z = -2.66 |
|  | MD = 105 | MD = 68.5 | p = 0.008* |
|  | SD = 47.52 | SD = 52.38 |  |
| **HQ** | M = 15.13 | M = 15.63 | z = 0.339 |
|  | MD = 15.5 | MD = 17 | p = 0.734 |
|  | SD = 7.53 | SD = 8.6 |  |
| **IHS** | M = 44.5 | M = 47 | z = -0.07 |
|  | MD = 43.5 | MD = 39.5 | p = 0.944 |
|  | SD = 17.67 | SD = 20.76 |  |

### Symptom Scores for Hyperacusis Questionnaires

Significant main effects of group were observed in cross-group comparisons between Acute, Chronic, and Controls for hearing loss in participants whose data were used in the analysis of response to at 1 kHz. Tukey’s post-hoc test revealed a non-significant trend between Acute Tinnitus and Chronic Tinnitus and between Acute Tinnitus and Controls, there were significant differences observed between Chronic Tinnitus and Controls. Controls had lower HQ scores compared to both Acute Tinnitus and Chronic Tinnitus. Consistent findings were obtained for IHS, and as shown by Tukey’s post hoc test, Controls had lower IHS scores in comparison to Acute Tinnitus and Chronic Tinnitus. There were statistical differences between Acute and Chronic Tinnitus too. Upon the paired longitudinal comparison, no change over time was observed for scores on either HQ or IHS.

For subjects included in the tinnitus frequency analysis, statistically significant equivalent results were obtained for both HQ and IHS across groups. Tukey’s post hoc test revealed non-significant trend differences between Acute Tinnitus and Chronic Tinnitus, significant differences between Acute Tinnitus and Controls, and significant differences between Chronic Tinnitus and Controls. Controls had lower HQ scores compared to both Acute and Chronic Tinnitus respectively. In comparison to Acute Tinnitus and Chronic Tinnitus, controls yielded lower IHS scores. There were significant differences noted between Acute Tinnitus and Chronic Tinnitus. Upon the paired longitudinal comparisons, no change over time was observed for scores on either HQ or IHS. The results of the symptom questionnaires and tinnitometry for each of the three groups are shown in table 2.

### ASSR Amplitude

The ASSR mean amplitude was compared between Acute, Chronic, and Controls across both frequencies and intensities with using 3-Way ANOVA. Results revealed significant main effect for group (F(2,574) = 8.008, p <0.001), significant main effect for frequencies (F(1,574) = 37.418, p <0.001), and significant main effect for intensities (F(2,574) = 9.83, p <0.001). With respect to differences in group, Controls (M = 1459.58) had increased ASSR amplitude when compared to both Acute Tinnitus (M = 1242.06) and Chronic Tinnitus group (M = 1108.09). However, no significant interaction effect for and for group, frequency, and intensity (F(4,573) = 0.479, p = 0.751) was obtained. Figure 3 depicts the differences in ASSR amplitude across group, frequency, and intensity. For the longitudinal comparison, a 3 way repeated measures ANOVA with a Greenhouse-Geisser correction determined that there was no interaction effect between group, frequency, and intensity (F(1.33,14.60) = 0.504, p = 0.541) on the ASSR ampltidue. Figure 4 depicts the differences in ASSR amplitude between Acute and Post Acute Tinnitus across frequency and intensity.

**Figure 3:**
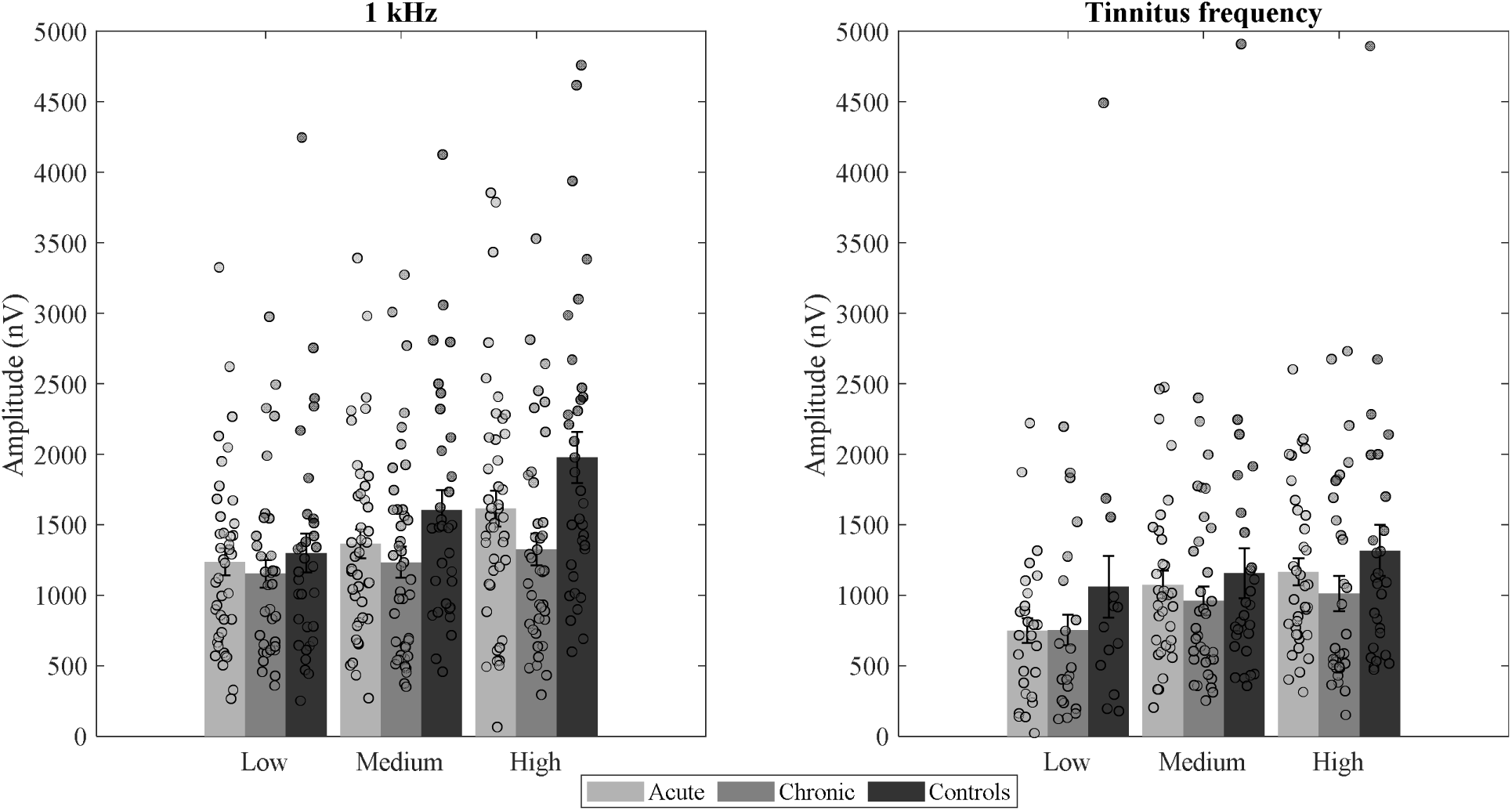
Differences in ASSR amplitude at low, medium, and high intensities between Acute, Chronic, and Controls for both 1 kHz and tinnitus frequency. There is no statistical difference between the groups for any intensities across either frequencies.

**Figure 4:**
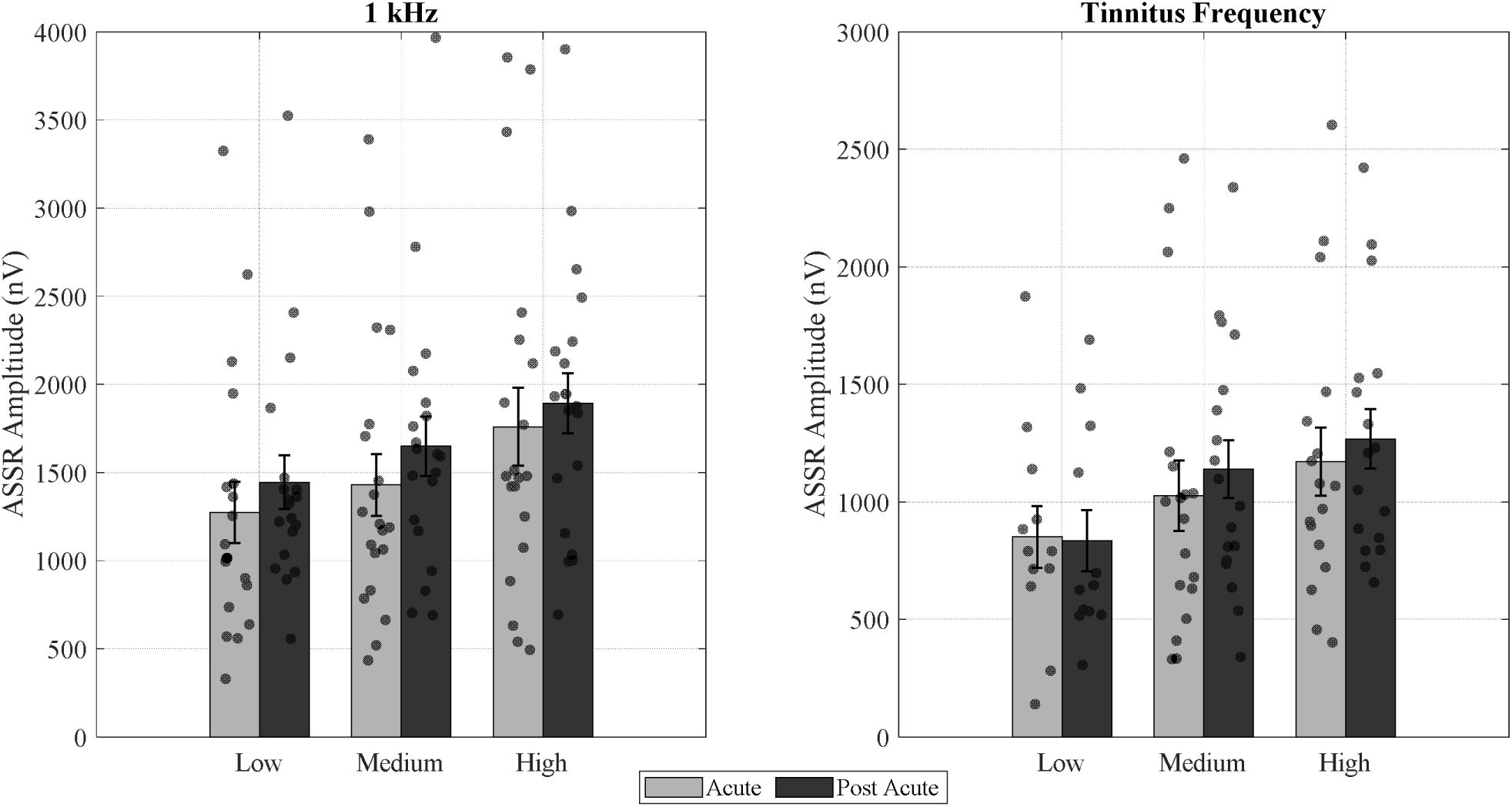
Differences in ASSR amplitude at low, medium, and high intensities between Acute and Post Acute Tinnitus for both 1 kHz and tinnitus frequency. There is no statistical difference between the groups for any intensities across either frequencies.

### ASSR Slope

In analysing the variations in slope among the Acute, Chronic, and Control groups, as well as between the Acute and Post Acute Tinnitus groups, we focused on three variables: the Stimulus Dynamic Range (SDR), defined as the difference between high and low stimulus presentation; the Amplitude Dynamic Range (ADR), which represents the difference between high and low ASSR amplitude; and slope, calculated as the ratio of SDR to ADR. Determining whether SDR differed will provide insight into the possible confounding factors that might affect the obtained ADR and slope. Assessing ADR in conjunction with slope will provide a comprehensive understanding of the amplitude growth function for ASSR across different groups.

### 1 kHz stimuli

A one-way MANOVA was carried out to establish the effect of group on slope, SDR, and ADR; this revealed a significant difference in overall response based on groups (F(6,184) = 3.964, p < 0.001, Wilk’s A = 0.526, partial η^2^ = 0.114). An overall main effect was seen for SDR (F (2,94) = 10.215, p < 0.001, partial η^2^ = 0.179) with difference between Acute Tinnitus and Controls (p = 0.007), Chronic Tinnitus and Controls (p < 0.001), and no differences between Acute Tinnitus and Chronic Tinnitus (p = 0.317). Controls had higher SDR (M = 19.74, SD = 5.94) when compared to Acute (M = 15.42, SD = 5.81) and Chronic Tinnitus groups (M = 13.19, SD = 4.95). There was an overall main effect of group seen for ADR (F (2,94) = 6.346, p = 0.003, partial η^2^ = 0.119) with difference between Acute Tinnitus and Controls (p = 0.049), difference between Chronic Tinnitus and Controls (p =0.002), and no difference between Acute Tinnitus and Chronic Tinnitus (p = 0.585). Controls (M = 640.39, SD = 396.70) had higher ADR when compared to Acute Tinnitus (M = 435.73, SD = 344.19) and Chronic Tinnitus (M = 328.49, SD = 263.19). There was, however, no main effect of groups on slope (F (2,94) = 0.961, p = 0.386, partial η^2^ = 0.02). There was an effect on SDR of group, which could also be a potential confound. This warranted us to carry out individual one-way ANCOVA to establish the effect of the group on ADR and slope individually, with covariates of SDR and hearing thresholds at 1 kHz (refer figure 5). With respect to Slope and ADR, there was no significant effect of group on the values of slope (F(2,92) = 1.279, p = 0.279, partial η^2^ = 0.027) and there was no significant effect of group on the values of ADR (F(2,92) = 1.774, p = 0.17, partial η^2^ = 0.037) (refer figure 5 and 6). Longitudinal comparisons between Acute Tinnitus and Post Acute Tinnitus revealed no significant difference in the slope values (t(17) = 0.806, p = 0.431), ADR (t(17) = 0.319, p = 0.754), and the SDR (t(17) = -1.177, p = 0.256) values respectively.

**Figure 5:**
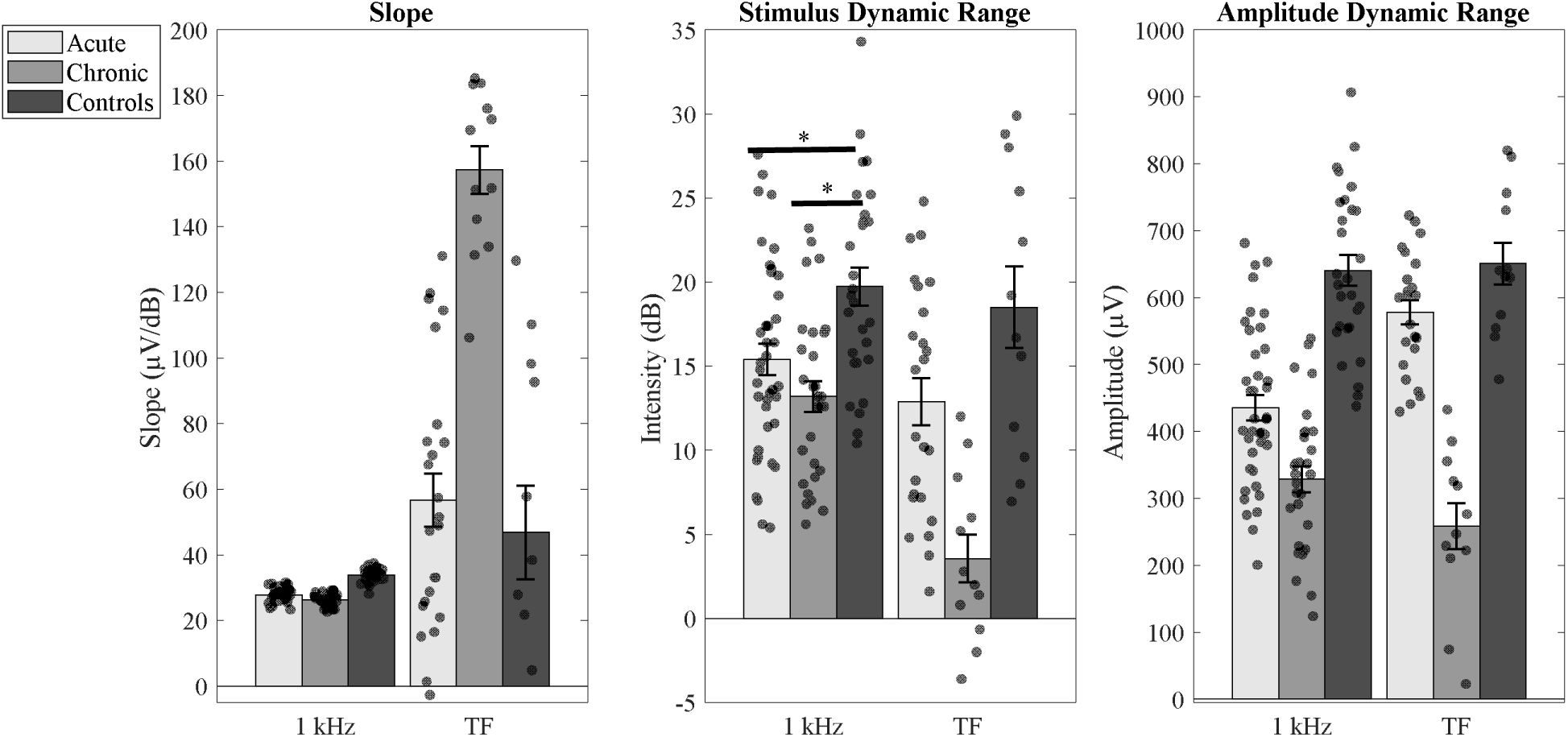
Slope, Stimulus Dynamic Range, and Amplitude Dynamic Range between Acute, Chronic, and Controls at both 1 kHz and tinnitus frequency. There is a significant difference between Acute Tinnitus and Controls and between Chronic Tinnitus and Controls for Stimulus Dynamic Range with controls having higher Stimulus Dynamic Range. No significant differences were found between other variables and groups.TF indicates Tinnitus Frequency and Asterix (*) indicates presence of significant difference

**Figure 6:**
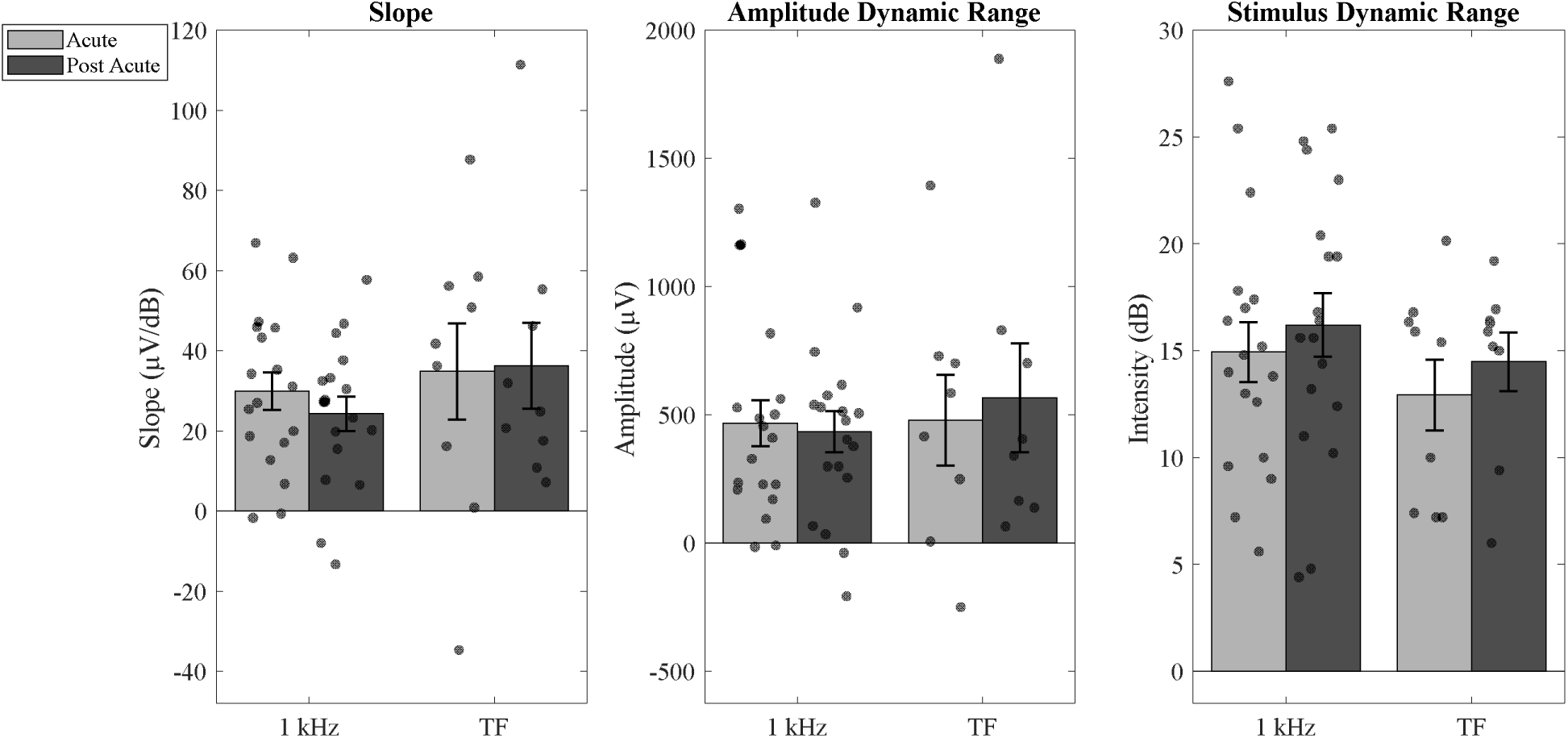
Slope, Stimulus Dynamic Range, and Amplitude Dynamic Range between Acute and Post Acute Tinnitus at both 1 kHz and tinnitus frequency. There were no significant differences in slope, stimulus dynamic range, and amplitude dynamic range between the groups across both frequencies. TF indicates Tinnitus Frequency

### Subjects included in tinnitus frequency

Similar analysis was carried out for the tinnitus frequency using non parametric Kruskal Wallis test to establish the effect of group on Slope, SDR, and ADR, and showed no significant effect on slope (*X^2^(2)* = 1.818, p = 0.4), non-significant trend for ADR (*X^2^(2)* = 5.286, p = 0.071), and presence of statistical significance for SDR ((*X^2^(2)* = 17.806, p < 0.001). Due to possibility of SDR being a confound for slope and ADR, we further ran a Quade non-parametric ANCOVA exploring the effects of groups on slope and ADR with SDR and hearing at 8 kHz as covariates. Results reveal no statistical differences for slope (*X^2^(2)* = 0.078, p = 0.925) and for ADR (*X^2^(2)* = 0.731, p = 0.487). Longitudinal comparisons between Acute Tinnitus and Post Acute Tinnitus reveals no significant difference in the slope values upon the Wilcoxon sign ranked test (*X^2^* (8) = -0.059, p = 0.953) and no significant difference in the SDR (*X^2^* (8) = -1.007, p = 0.314) values too between the groups (refer figures 5 and 6).

## Discussion

The principal results indicate that there were no significant differences longitudinally between Acute and Post Acute Tinnitus as well as cross sectionally between Acute, Chronic, and Controls for both ASSR ampltidue and slope. We did however get a main effect of group with Controls having higher ASSR amplitude when compared to both Acute and Chronic Tinnitus.

### Results are not confounded by extraneous factors

Although we did not yield pure statistically significant results, it is essential to emphasise that the potential extraneous or confounding factors have either been controlled or did not impact the current results. The current study had groups that were controlled for age, sex, and stimulus ear presentation. The degree of hyperacusis differed between the groups, and we have adjusted the individual’s dynamic range for recruitment, hyperacusis/gain, and to ensure the stimulus was delivered within the participant’s comfortable range. Our results were repeated using a robust longitudinal pairwise comparison design between Acute and Post Acute stages, giving these findings additional robustness to any remaining potential confounding factors in the cross-sectional comparison between Acute and Chronic Tinnitus.

### Gross increase of ASSR in controls

Our initial findings indicate that there were no cross-sectional changes between the Acute, Chronic, and Control groups, with respect to either slope and absolute amplitude across freqeuncies and inteisities. We however report grossly increased ASSR amplitude in Controls when compared to Acute and Chronic through main effects of groups on ASSR amplitude.

Supporting literature highlight the role of attentional modulation on the ASSR ampltidue. Roberts et al. (2013), explored the role of attention and tinnitus with respect to ASSR and they observed a smaller ASSR responses in individuals with tinnitus subjects when compared to the Controls at specific frequencies. A study by Paul et al. (2014) demonstrated that individuals with tinnitus exhibit impaired attentional modulation of the 40 Hz auditory steady-state response specifically within their tinnitus frequency range. The controls tend to show increased ASSR amplitude with attention at both 500 Hz and 5 kHz, but the tinnitus individuals tend to show attention modulation at 500 Hz (below their tinnitus frequency) and not 5 kHz.

Other studies highlight the potential role of distress on the ASSR amplitude, for instance Sadeghijam et al. (2022) suggested a relationship between tinnitus distress, attention networks, and ASSR amplitudes. Specifically, they propose that tinnitus individuals with high distress tend to have an increased activity around the prefrontal regions (which is strongly associated with attention and distress networks) and thereby leading to lower ASSR amplitudes. The reduction in ASSR amplitudes can also be due to the overlap between these attention and distress networks with the ASSR generating networks, potentially causing interference. On the other hand, those with lower tinnitus distress may exhibit less prefrontal involvement and thereby a higher ASSR amplitudes due to reduced inhibition or reduced overlapping between the distress networks and ASSR generating network. The article also further highlights that the generation of an ASSR and the robustness of it is based on the combination of the activity of auditory and prefrontal network regions, and individuals with Chronic Tinnitus with severe distress tend have abnormal ASSR amplitude due to inhibitory dysfunction. The article by Moossavi et al. (2019) explores the relationship between Auditory Steady-State Response amplitudes and tinnitus distress levels through a brief pilot study. The levels of tinnitus distress were divided based on the THI scores: a high-stress group and a low-stress group. The researchers measured the amplitudes of the 40 Hz ASSR in both the stress groups and their findings reveal that individuals that individuals with low distress tend to have increased ASSR when compared to individuals with high distress.

In our result, both abnromal attention and presence of distress amongst the tinnitus participants may have contributed to the reduction of ASSR amplitude when compared to controls.

We could also infer reduction of ASSR in individuals with tinnitus to be attributed to desynchronization caused by the tinnitus. A study by Motomura et al. (2024) investigated the impact of abrupt sound pressure changes on the 40-Hz auditory steady-state response.

Transient frequency analysis was employed to examine the relationship between the magnitude of sound pressure change and both the amplitude and inter-trial phase coherence of the 40-Hz ASSR. The ASSR was elicited by a click train with a fixed inter-click interval, and an abrupt change in sound pressure was introduced midway through the train. The results demonstrated that the abrupt change caused desynchronization of the 40-Hz ASSR, and the degree of desynchronization was dependent on the magnitude of the sound pressure change. In our case, we attribute that the tinnitus is an ongoing neural response that increases the desynchronization causing reduction of the magnitude of ASSR.

Lastly, the role of GABA (Gamma-Aminobutyric acid) has been linked to changes in ASSR amplitude. Sugiyama et al. (2021) found that boosting GABAergic transmission via lorazepam, a benzodiazepine, increased the 40 Hz ASSR in the early auditory cortex. Another study study by Toso et al. (2024) examined the roles of GABAergic and NMDA receptor-mediated synaptic interactions in generating the 40 Hz auditory steady-state response (ASSR) in the human auditory cortex. Using MEG and placebo-controlled pharmacological interventions, healthy participants received lorazepam (a GABAA_AA receptor agonist) and memantine (an NMDA receptor antagonist). Lorazepam significantly increased 40 Hz ASSR amplitude, indicating a key role for GABAergic inhibition in modulating this response. In contrast, memantine had no effect, suggesting NMDA receptor-mediated excitation may be less critical. These findings support the use of 40 Hz ASSR as a non-invasive marker of cortical circuit function, relevant to disorders like schizophrenia. With respect ot GABA and tinnitus, a prevailing theory is that the reduced GABAergic inhibition in the auditory pathway may contribute to the onset of tinnitus due to increase in hyperexcitability and this has reflected with the presence of lower levels of GABA at the level of the auditory cortex for tinnitus individuals (Sedley et al., 2015; Isler et al., 2022). As Toso et al. (2024), emphasized the association between GABAergic activity and ASSR amplitude variations, the reduced ASSR amplitudes observed during tinnitus condition in our study may suggest a potential link with GABAergic mechanisms considering there were reduced ASSR amplitudes which could be framed as hypotheses that might inform future studies through GABA-sensitive techniques such as magnetic resonance spectroscopy (MRS) (Sedley et al., 2015).

### No changes in sensitivity between Acute and Chronic Tinnitus

Despite gross changes in amplitude between Controls and Acute/Chronic Tinnitus, we yielded no statistical differences between Acute and Post Acute Tinnitus or between Acute and Chronic Tinnitus. We infer that despite fluctuations in tinnitus activity, characterised by a reduction in tinnitus loudness and distress over time (see tables 2 and 3), neural sensitivity remains consistent across groups, demonstrating that tinnitus activity is independent of auditory sensitivity. This is consistent with our recent work (Umashankar, Gander, et al., 2025), which found that despite a reduction in tinnitus loudness and distress over time, subject sound sensitvity did not change between the groups, indicating that the level of hyperacusis within the tinnitus is what causes these alterations. These findings contrast with our previous research, which indicates heightened hypersensitivity in Acute Tinnitus compared to Post Acute Tinnitus (Umashankar, Sedley, et al., 2025; Umashankar, Alter, et al., 2025). We conclude that both neural sensitvity and subjective auditory sensitvity remain constant, indicating whether central gain is actually associated to tinnitus.

### Summary

The current study indicates that grossly increased ASSR amplitudes in controls is possibly impacted by alterations in neural synchronisation, tinnitus-related distress, GABAergic inhibition and/or attentional modulation. Howver, there were no variations with the ASSR amplitude between Acute and Chronic Tinnitus or even between Acute and Post Acute Tinnitus which possibly suggests that the tinnitus activity as such might be indendpent from neural sensitvity measures like central gain. Figure 7 is a hypothetical model that integrates the above-mentioned highlighting the presence of tinnitus as a continuous stimulation that diminishes phase locking to the incoming signal, while no tinnitus enhances phase locking to the incoming signal.

**Figure 7:**
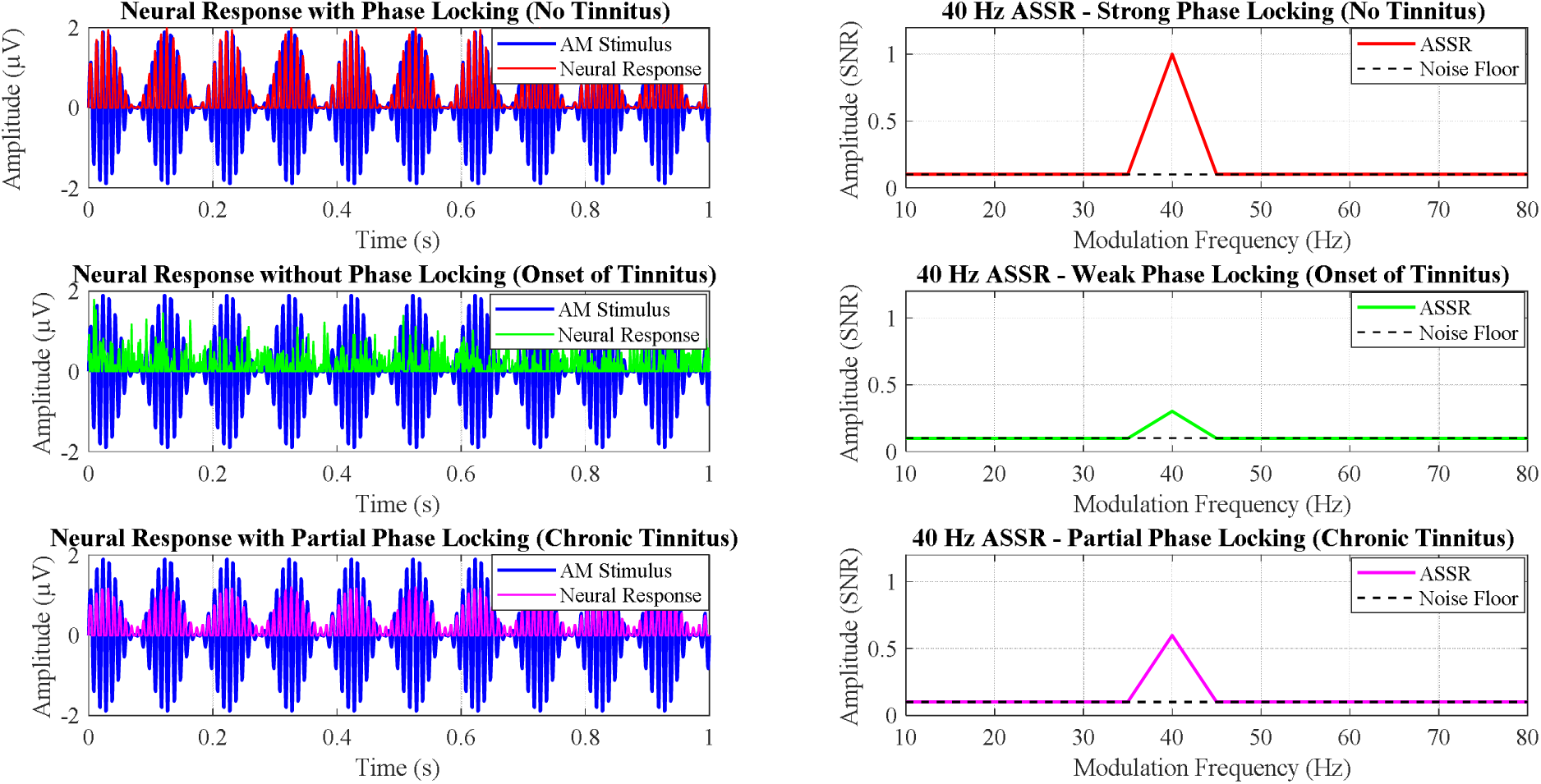
Differences in neural response of AM tones across 3 conditions of No Tinnitus, Acute Tinnitus, and Chronic Tinnitus. In the absence of tinnitus, there exists sufficient neuronal synchrony (phase locking), allowing the neurons to respond appropriately to the amplitude-modulated stimuli. The onset of tinnitus diminishes neural synchronisation as the neurons that should respond to the amplitude-modulated tones now produce a continuous ongoing stimulation (spontaneous activity), hence decreasing phase locking to the incoming signal. In the chronic stages of tinnitus, the tinnitus activity diminishes, leading to a partial reduction in abnormal neural synchronisation and an enhancement of the auditory steady-state response (ASSR) amplitude moving towards controls.

